# Mapping a mutation causing pale yellow petals in *Brassica rapa*

**DOI:** 10.1101/2022.10.17.512571

**Authors:** Hiu Tung Chow, Rebecca A. Mosher

## Abstract

Petal color is an important trait for both ornamental purposes and also for attracting pollinators. Here, we report a mutation of *Brassica rapa* R-o-18 with pale yellow petals that we retrieved from an EMS population and named *whiter shade of pale* (*wsp*). Phenotypic segregation ratio of an F2 mapping population indicates the phenotype is controlled by a single recessive gene. Mapping data from the whole genome sequencing coupled with allele frequency analysis suggests the mutation is located in a ∼2 Mbp interval on chromosome 2. The interval contains a putative esterase/lipase/thioesterase protein previously demonstrated to account for floral color in *B. rapa*. We demonstrate that *wsp* carries a G to A missense mutation causing an aspartate to asparagine substitution within the putative lysophospholipid acyltransferase domain.

## Description

Petals are often brightly colored to attract insects for pollination (Maharaj et al., 2021). Petal color is controlled by pigments such as carotenoids, which are synthesized and stored in organelles called chromoplasts (Zhu et al., 2010; Sobel and Streisfeld, 2013). Mutations of carotenoid biosynthesis enzymes can alter the accumulation of different carotenoids, resulting varying hues of yellow, orange, or red (Zhu et al., 2010). Some species, such as *Arabidopsis thaliana*, presumably lack chromoplasts in their petal cells and thus produce white petals without carotenoid accumulation (Pyke and Page, 1998).

While the majority of *Brassica rapa* varieties have bold yellow flowers, some breeding lines display pastel yellow petals (Zhang et al., 2020; Yang et al., 2021; Rahman, 2001). Mutations responsible for this phenotype were cloned, uncovering enzymes impacting carotenoid accumulation. One such mutation is *BrWF3*, a homolog of Arabidopsis *PHYTYL ESTER SYNTHASE 2*, which encodes a putative acyltransferases in the esterase/lipase/thioesterase family. A double-haploid line with a frame shift mutation at *BrWF3* displays pale white petals and reduced carotenoid accumulation (Yang et al., 2021). The frame shift causes truncation of the protein and loss of conserved enzymatic domains (Yang et al., 2021). Similarly, genetic control of white flowers in an ornamental *Brassica juncea* line was mapped to two loci containing the homeologs of *BrWF3* (*BjuB027334* and *BjuA008486*) (Zhang et al., 2018a, 2018b). In white flowered individuals, *BjuB027334* carrys a missense mutation and has significantly decreased mRNA accumulation, while the genetic change at *BjuA008486* is unknown.

Here, we identified a mutant with pale yellow petals from an EMS mutagenized population of *B. rapa* variety R-o-18 (**Figure 1A**). We named this mutant *whiter shade of pale* (*wsp*) given its pale yellow petals. To map the mutation, we crossed a single *wsp* individual with *B. rapa* variety R500, which carries numerous single nucleotide polymorphisms (SNPs) that can be used for genetic markers during mapping (**Figure 1B**). We propagated the cross to the F2 generation and grew 718 F2 individuals, of which 183 individuals (25.5%) exhibited the *wsp* phenotype. Coupled with the fact that all F1 plants possessed bright yellow petals, the ∼ 3:1 phenotypic segregation of the wild-type to *wsp* phenotype in the F2 population indicates that the phenotype is controlled by a recessive mutation (p-value = 0.09, chi-square test). DNA from the 183 F2s with mutant phenotype was pooled equally and prepared for whole genome sequencing. By calculating allele frequency of SNPs covered by sequence reads, we mapped the mutation to ∼2 Mbp on chromosome 2 (**Figure 1C**). Next, we annotated EMS-derived mutations (C to T or G to A) within the mapping interval, resulting in 105 sites with quality score >30. Four sites were predicted as missense mutations, 5 were predicted as silent mutations, and 96 fell in predicted noncoding regions. The 4 genes harboring missense mutations are not known to be involved in carotenoid or chromoplast synthesis, however *BrWF3* is also located within the mapping interval and contains a predicted intronic mutation (quality score = 228). Because an intronic mutation is unlikely to disrupt enzyme function, we suspected that the gene was incorrectly annotated in the R-o-18 genome assembly. Comparison of this annotation (*A02p049750*.*1_BrROA*) to the published sequence or with *in silico* predictions of coding regions confirmed this suspicion (Yang et al., 2021). The original intronic mutation falls within the reannotated transcript, causing a G to A mutation at the 1729^th^ nucleotide and replacing an aspartic acid with asparagine at the 577^th^ amino acid (**Figure 1D**). We predicted the protein structure using AlphaFold2 and also annotated the mutation using Missense3D (Ittisoponpisan et al., 2019; Jumper et al., 2021). The substitution is predicted to cause structural damage due to substituting a buried charged residue with an uncharged residue. The loss of negative charge of the aspartic acid might induce structural instability of the enzyme. Alternatively, since *wsp* phenocopies presumed null alleles of *BrWF3*, this mutation, which falls within the highly conserved putative lysophospholipid acyltransferases domain (**Figure 1F**), might prevent the transfer of acyl groups to substrates.

**Figure 1.**
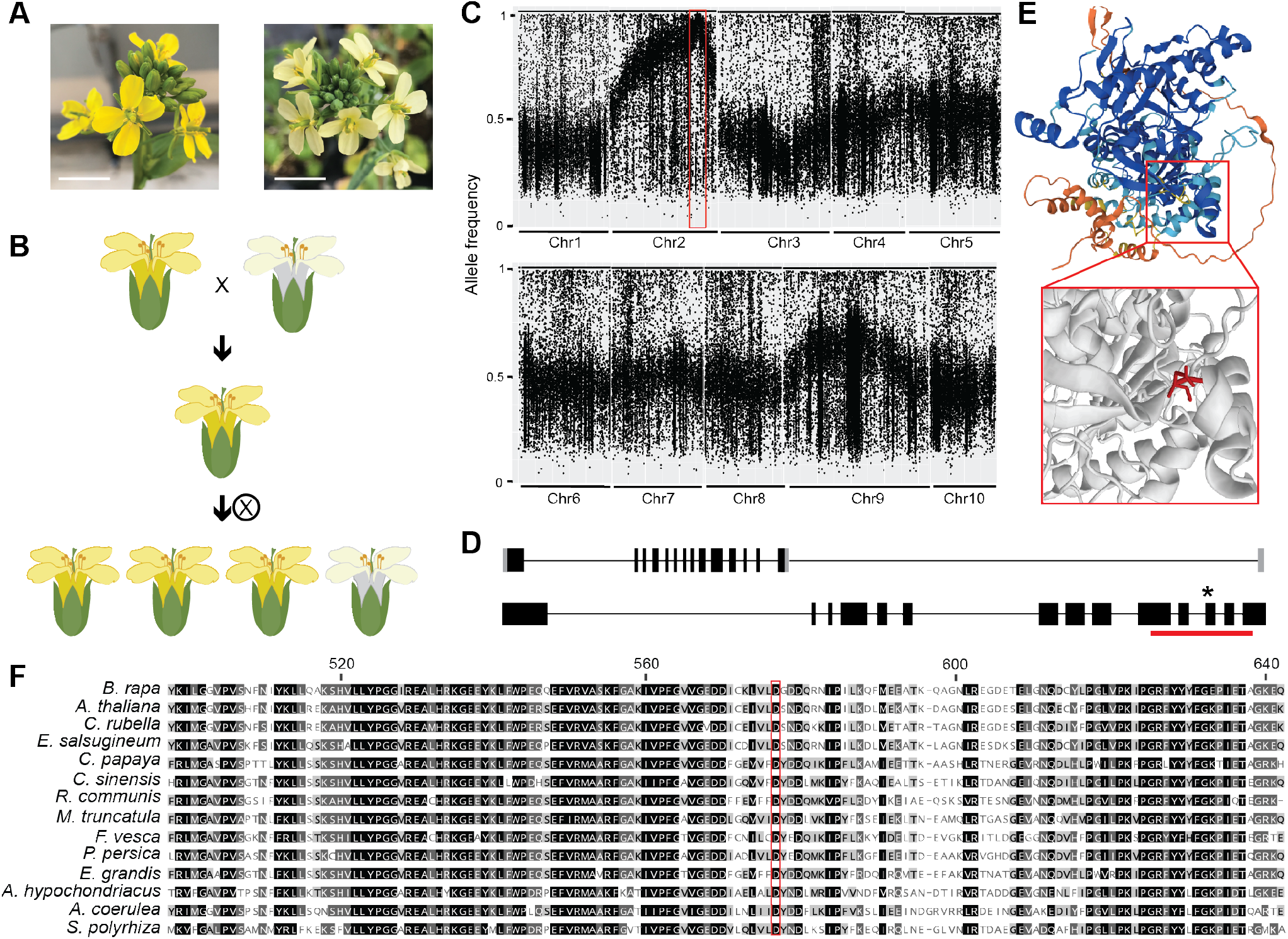
*whiter shade of pale*, a pale yellow petal *Brassica rapa* mutant, carries a mutation in *BrWF3*. ***(A)*** *B. rapa* R-o-18 WT (left) has bright yellow flower, while *wsp* petals (right) are pale yellow. Scale bar = 1cm. **(B)** Crossing scheme to generate the F2 mapping population. *wsp*, which is from R-o-18 background, was crossed to R500 wild type to generate an F1 with yellow petals. The F1 was self-crossed to generate the F2 generation, in which 25% of individuals had pale petals. **(C)** Allele frequencies of R-o-18 by R500 SNP variants over chromosomes among pooled pale yellow individuals from the F2 population. Red box indicates the ∼ 2Mbp mapping interval on chromosome 2. **(D)** Gene structure of original (top) and improved (bottom) annotations of *A02p049750*.*1_BrROA*. Black lines indicate introns while black bars indicate exons; untranslated regions in the original annotation are represented by gray boxes. An asterisk denotes the *wsp* missense mutation and the orange line indicates the region encoding the putative lysophospholipid acyltransferase domain. **(E)** Predicted protein structure of A02p049750.1_BrROA (top) colored by pLDDT confidence measure on AlphaFold. The boxed region is magnified below, with the site of the *wsp* mutation labelled red (bottom). **(F)** Alignment of the partial amino acid sequence of *A02p049750*.*1_BrROA* with closest homologs from other plant species. Comparison of A02p049750.1_BrROA with *Arabidopsis thaliana* (*A. thaliana*; At3g26840.1), *Capsella rubella* (*C. rubella*; Carub.005s0494.1), *Eutrema salsugineum* (*E. salsugineum*; Thhalv10003736m), *Carica papaya* (*C. papaya*; evm.model.supercontig_70.97), *Camellia sinensis* (*C. sinensis*; orange1.1g041641m), *Ricinus communis* (*R. communis*; 30131.m007010), *Medicago truncatula* (*M. truncatula*; Medtr7g083200.1), *Fragaria vesca* (*F. vesca*; FvH4_3g35400.t1), *Prunus persica* (*P. persica*; Prupe.3G311200.1), *Eucalyptus grandis* (*E. grandis*; Eucgr.101613.1), *Amaranthus hypochondriacus* (*A. hypochondriacus*; AH010860-RA), *Aquilegia coerulea* (*A. coerulea*; Aqcoe5G342400.1), and *Spirodela polyrhiza* (*S. polyrhiza*; Spipo6G0003600). Identical amino acid residues are rendered black. Similar residues in at least 80% all comparing species are highlighted in dark-grey, whereas similar residues in at least 60% but lower than 80% similarity are highlighted in light-grey. The red box marks the mutated residue in *wsp*. Numbers along the top refer to the amino acid position within A02p049750.1_BrROA.

This work confirms that *A02p049750*.*1_BrROA* (*BrWF3*) is involved in petal coloration in *Brassica rapa*, including in the R-o-18 variety. The allele we describe also provides genetic resource for functional characterization of this enzyme in carotenoid accumulation.

## Methods

### Plant materials, growth condition and sequencing

All plants were grown in a greenhouse at 18°C with 16 hour supplemental lighting. The *wsp* mutation was created in the R-o-18 variety and the R500 variety was used for mapping. Ethyl methanesulfonate (EMS) treatment was performed as described in (Chow et al., 2022) and *wsp* was recovered in the M2 generation. Bulked Segregant Analysis sequencing was performed following (Chow et al., 2022). The reannotated protein sequence was deposited in NCBI GeneBank (OP610629). Sequencing data from this article can be retrieved in NCBI SRA under accession number PRJNA888995.

### Prediction of impact of variant on protein structure

The A02p049750.1_BrROA protein sequence was used as a query at Phytozome to recover homologous sequences shown in Figure 1F and also at UniProt (www.uniprot.com). Entry M4EVX4_BRARP from the *B. rapa* variety Chiifu is 100% identical and its predicted protein structure was obtained from AlphaFold (Jumper et al., 2021). The impact of the *wsp* mutation on protein structure was predicted using Missense3D (Ittisoponpisan et al., 2019).

## Acknowledgements and Funding

This work is supported by project number ARZT-3039860-G25-578 from the USDA National Institute of Food and Agriculture to RAM. DNA sequencing was conducted by the University of Arizona Genetics Core.

## Notes

### Competing Interest Statement

The authors have declared no competing interest.

## References

Chow, H.T., Kendall, T., and Mosher, R.A. (2022). A novel CLAVATA1 mutation causes multilocularity in Brassica rapa. bioRxiv.

Ittisoponpisan, S., Islam, S.A., Khanna, T., Alhuzimi, E., David, A., and Sternberg, M.J.E. (2019). Can predicted protein 3D structures provide reliable insights into whether missense variants are disease associated? J. Mol. Biol. 431: 2197–2212.

Jumper, J. et al. (2021). Highly accurate protein structure prediction with AlphaFold. Nature 596: 583–589.

Maharaj, G., Bourne, G., and Ansari, A. (2021). A review of floral color signals and their Heliconiid butterfly receivers. In Arthropods - Are They Beneficial for Mankind? (IntechOpen).

Pyke, K.A. and Page, A.M. (1998). Plastid ontogeny during petal development in Arabidopsis. Plant Physiol. 116: 797–803.

Rahman, M.H. (2001). Inheritance of petal colour and its independent segregation from seed colour in Brassica rapa. Plant Breed. 120: 197–200.

Sobel, J.M. and Streisfeld, M.A. (2013). Flower color as a model system for studies of plant evo-devo. Front. Plant Sci. 4: 321.

Yang, S., Tian, X., Wang, Z., Wei, X., Zhao, Y., Su, H., Zhao, X., Tian, B., Yuan, Y., and Zhang, X.-W. (2021). Fine Mapping and Candidate Gene Identification of a White Flower Gene BrWF3 in Chinese Cabbage (Brassica rapa L. ssp. pekinensis). Front. Plant Sci. 12: 646222.

Zhang, N., Chen, L., Ma, S., Wang, R., He, Q., Tian, M., and Zhang, L. (2020). Fine mapping and candidate gene analysis of the white flower gene Brwf in Chinese cabbage (Brassica rapa L.). Sci. Rep. 10: 6080.

Zhang, X., Li, R., Chen, L., Niu, S., Chen, L., Gao, J., Wen, J., Yi, B., Ma, C., Tu, J., Fu, T., and Shen, J. (2018a). Fine-mapping and candidate gene analysis of the Brassica juncea white-flowered mutant Bjpc2 using the whole-genome resequencing. Mol. Genet. Genomics 293: 359–370.

Zhang, X., Li, R., Chen, L., Niu, S., Li, Q., Xu, K., Wen, J., Yi, B., Ma, C., Tu, J., Fu, T., and Shen, J. (2018b). Inheritance and gene mapping of the white flower trait in Brassica juncea. Mol. Breed. 38.

Zhu, C., Bai, C., Sanahuja, G., Yuan, D., Farré, G., Naqvi, S., Shi, L., Capell, T., and Christou, P. (2010). The regulation of carotenoid pigmentation in flowers. Arch. Biochem. Biophys. 504: 132–141.

